# First record of *Venturia canescens* GRAVENHORST, 1829 (Hymenoptera: Ichneumonidae: Campopleginae) from Tunisia

**DOI:** 10.1101/2024.07.25.605196

**Authors:** Asma Zeiri

**Affiliations:** University of Carthage, Faculty of Sciences of Bizerte (FSB), Bizerte, Tunisia

**Keywords:** Endoparasitoid, Hymenoptera, Ichneumonidae, Tunisia, Venturia canescens

## Abstract

**Background:** *Venturia canescens* Gravenhorst 1829 (Hymenoptera: Ichneumonidae: Campopleginae), a solitary koinobiont endoparasitoid, lives on larvae of various Lepidopteran hosts belonging to the families of Pyralidae, Noctuidae, Tortricidae, Gelechiidae, Tineidae and Yponomeutidae of Lepidoptera (typically Pyralidae). It was frequently reported on the carob moth *Ectomyelois* ceratoniae (Zeller) (Lepidoptera: Pyralidae). The *E. ceratoniae* is an insect pest that causes damage to various crops and fruit trees throughout the world. It was recorded on dates *(Phoenix dactylifera)*, pomegranates (*Punica granatum*), almonds (*Prunus dulcis*) and pistachio nut (*Pistacia vera*), affecting the fruit quality. In Tunisia, this pest was recorded on pomegranate by many authors.

**Results:** The parasitoid *V. canescens*, is accidently recorded from Tunisia within the collection of parasitoids of the almond bark beetle *Scolytus amygdali* (Coleoptera: Curculionidae: Scolytinae) in the orchard of the Professional Training Center of Agriculture, Jammel, Monastir, Tunisia (35°37′60′′ N:10°46′0′′E) rich of fruit trees. The identification of *V. canescens* was carried out using key for females of the genus Venturia to Western Palaearctic species and the specimen is reported as a first record in Tunisia. No male was collected, the host is unknown.

**Conclusions:** This species is widely distributed in the Palaearctic region. As far as we know this is a first record for the country.

## Background

Worldwide 144 valid species of *Venturia* Schrottky, 1902 (Ichneumonidae, Campopleginae) have been reported (Han et al., 2021), of which 11 species were recorded in the Afrotropical region and 8 in the Western Palaearctic region (Cameron 1906; Marchesini and Dalla Monta 1994; Thiéry et al. 2001; Yu et al. 2012; Zwakhals and Van Achterberg 2017; Vas 2020; Vas and Di Giovanni 2020). An identification key to the Western Palaearctic species of *Venturia* was recently provided (Vas 2020; Rousse and Villemant 2012).

Among the genera of Campopleginae, the genus *Venturia* includes species reported as important for the control of agricultural pests and considered a cosmopolitan species, with a worldwide distribution and most often found in buildings where grains or flour are stored (Fig. 1A and B.) (Scaramozzino et al. 2018). Solitary larval parasitoid *Venturia canescens* (Gravenhorst 1829) (Hymenoptera: Ichneumonidae), is an endoparasitoid of several lepidopterous larvae belonging to the family Pyralidae (Scaramozzino et al. 2018). Its host spectrum contains many moth species. Known hosts from Pyralidae are: *Ephestia kuehniella* (Zeller 1879), *Plodia interpunctella* (Hübner 1813), *Apomyelois ceratoniae* (Zeller 1839), *Galleria mellonella* (Linnaeus 1758) *and Ostrinia nubilalis* (Hübner 1796); Tineidae: *Nemapogon granella* (Linnaeus 1758); Gelechiidae: *Phthorimaea operculella* (Zeller 1873); Yponomeutidae: *Prays citri* (Millière 1873); Tortricidae: *Grapholita funebrana* (Treitschke 1835) and some Noctuidae (Yu et al. 2012; Scaramozzino et al. 2018).

**Fig. 1.**
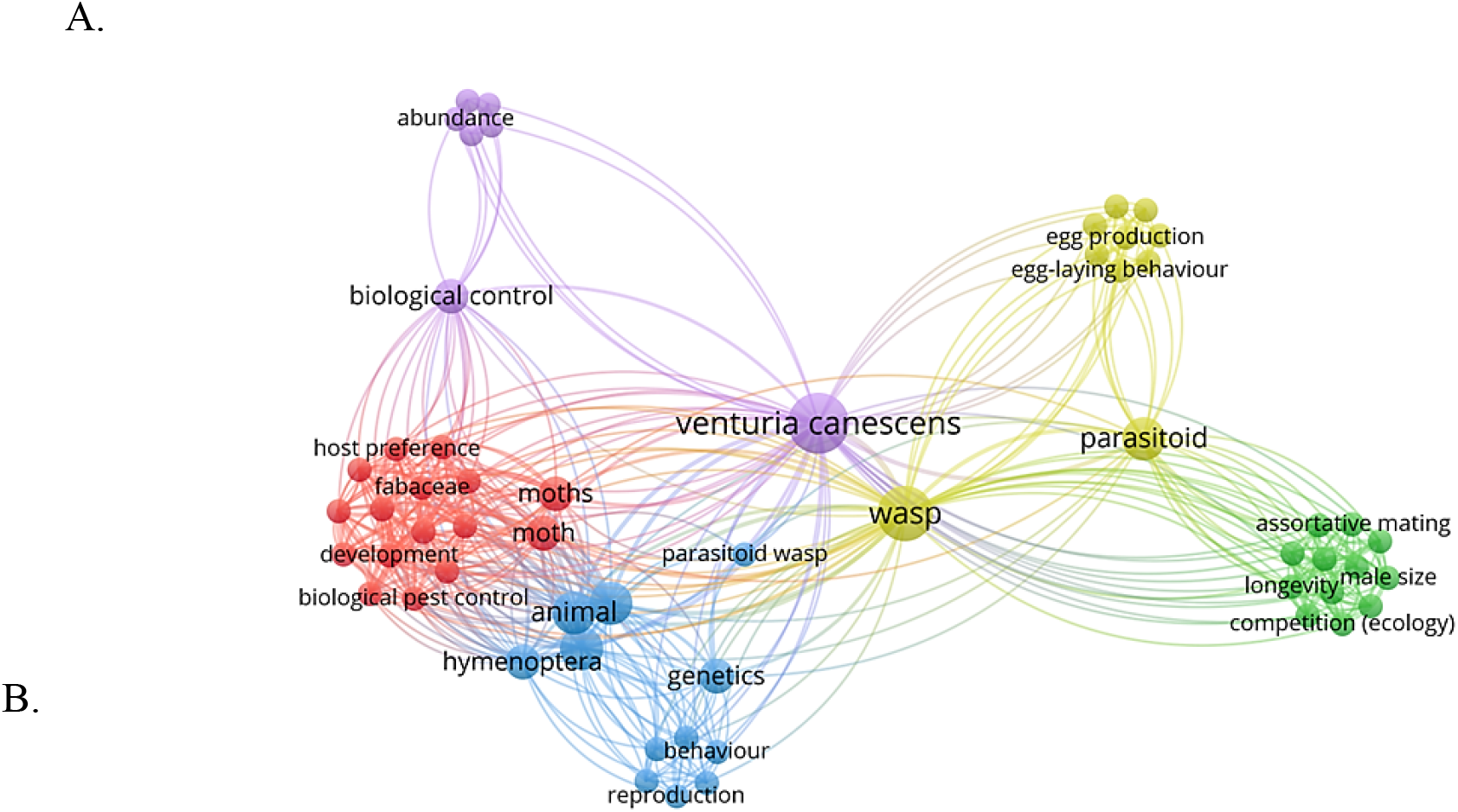
**A and B**. A network map representing the bibliometric analysis of scientific research in relation with *Venturia canescens*. This figure was generated using VOS Viewer (van Eck and Waltman, 2013), and the data utilized in this study were collected from Scopus database using following keywords: A: Venturia AND canescens OR pyralidae OR Noctuidae OR Tortricidae OR Gelechiidae OR Tineidae OR Yponomeutidae; B. Venturia AND canescens OR Africa OR Europa OR America OR Asia.

It is considered an occasional parasitoid of *Lobesia botrana*, of rather marginal importance (Marchesini and Dalla Monta 1994; Scaramozzino et al. 2018; Villemant et al. 2011). This species is very common and has a very wide geographical distribution. It has been repeatedly described under different names in combination with different genera (Yu et al. 2012; Wahl 1987; Yu and Horstmann 1997).

In this paper a first record of *V. canescens* from Tunisia is collected and identified.

## Methods

The material for this paper was accidentally obtained during the collection of parasitoids *Scolytus amygdali* Guérin-Méneville, 1847 in Tunisia. A female was collected in 2011 from the Sahel region of Tunisia (**Fig. 2**).

**Fig. 2.**
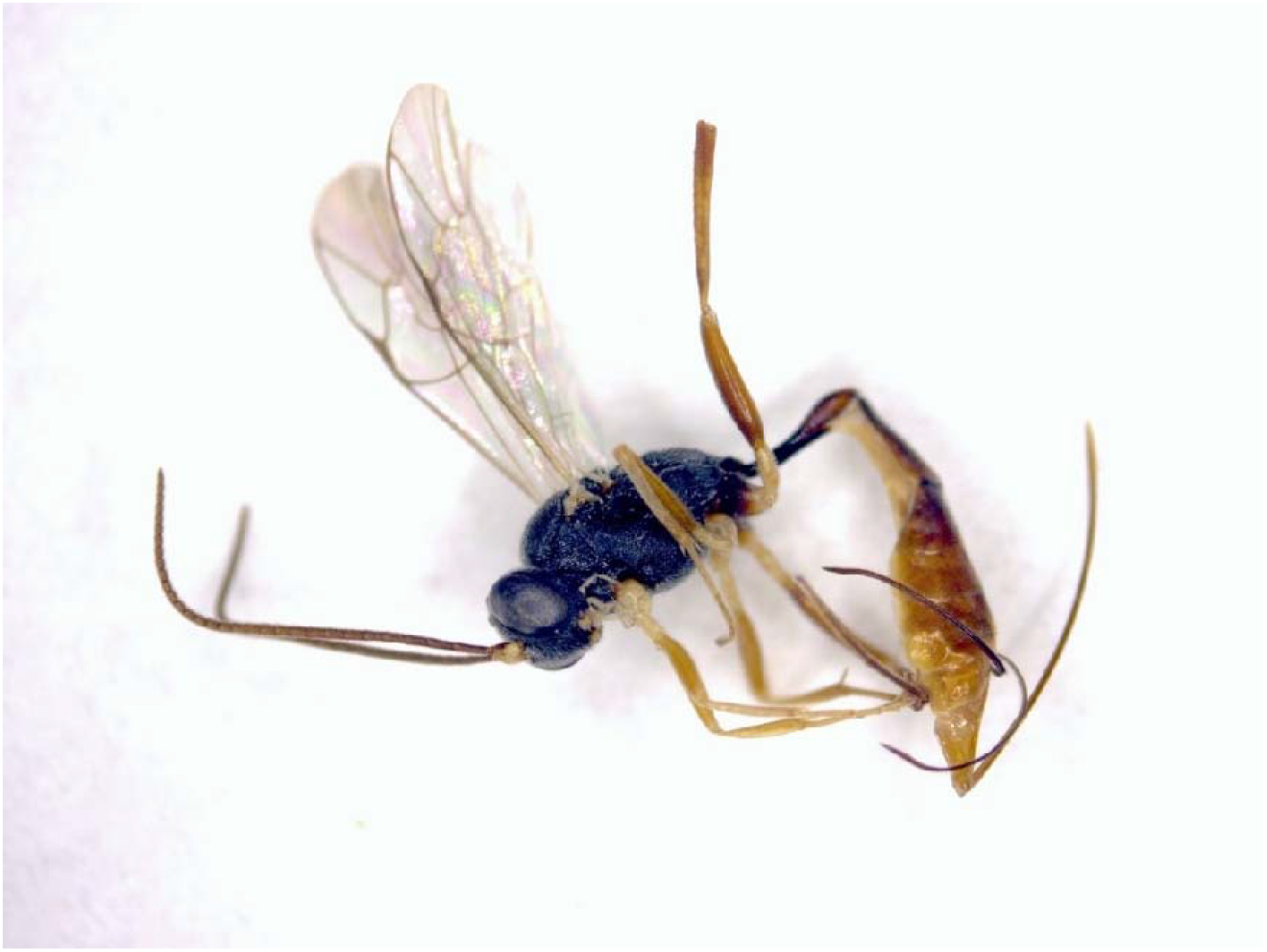
Female of *Venturia crescens* collected from the Sahel region of Tunisia.

The specimens were identified and examined by the author using a Carl Zeiss Stemi 508 stereoscopic microscope. Photos were taken with a Zeiss Axiocam 105 Color camera. The identification was carried out following keys published by Vas (2020) and Rousse and Villemant (2012). Some consulted reference images were digital. Distributional records of species were checked in Yu *et al*. (2012) and Van Noort (2022).

## Results

### Material examined (Fig. 3)

– 1♀, [Tunisia, 2011] leg. Zeiri, A., det. Zeiri, A.; specimen in alcohol. The male sex is unknown in the majority of the Western Palaearctic species. Host unknown.

**Fig. 3.**
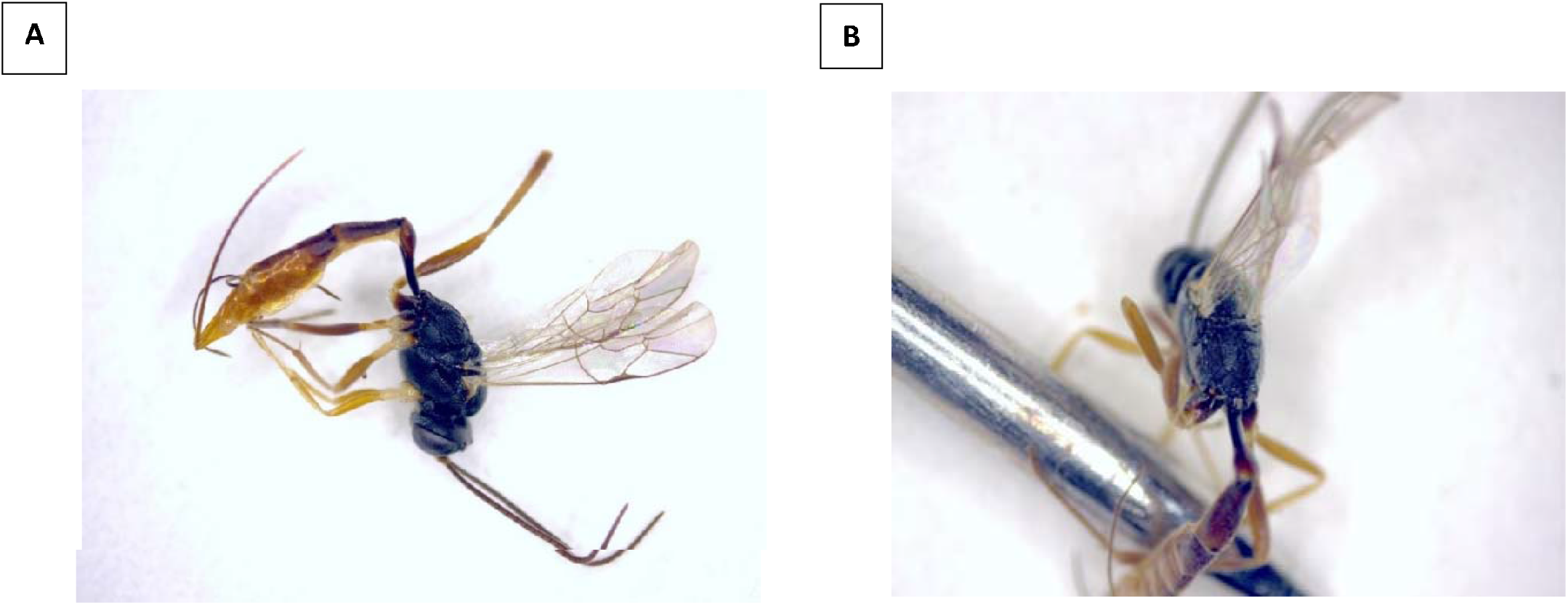
A Lateral and B. dorsal View of female of *Venturia crescens*

Subfamily: Campopleginae (Förster 1869)

Genus: *Venturia* (Schrottky 1902)

List of synonymies:

*Venturia canescens* (Gravenhorst 1829)

*Campoplex canescens* (Gravenhorst 1829)

*Devorgilla canescens* (Gravenhorst 1829)

*Nemeritis canescens* (Gravenhorst 1829)

*Venturia australica* (Girault 1925)

*Venturia christianae* (Cheesman 1928)

*Venturia compressa* (Hedwig 1962)

*Venturia ductilis* (Say 1835)

*Venturia ephestia (*Cameron 1912)

*Venturia ephestiae* (Cameron 1912)

*Venturia frumentaria* (Rondani 1874)

*Venturia garrula* (Cameron 1905)

*Venturia gracilens* (Tosquinet 1896)

*Venturia insularis* (Ashmead 1901)

*Venturia oahuensis* (Ashmead 1901)

*Venturia orientalis* (Schmiedeknecht 1909)

*Venturia pyraustae* (Uchida 1930)

### Diagnosis (Fig. 4)

Mesopleuron distinctly punctate; metapleuron and propodeum punctate; Forewing with areolet; areolet of forewing not triangular; Pterostigma brown, superomedial area pentagonal and distinctly elongated. Hind femur not black; Metasoma predominantly dark /reddish. Ovipositor sheath at most 2× (usually 1.5–1.6×) as long as hind tibia. Clypeus in profile flat. Postpetiolus is reddish brown.

**Fig. 4.**
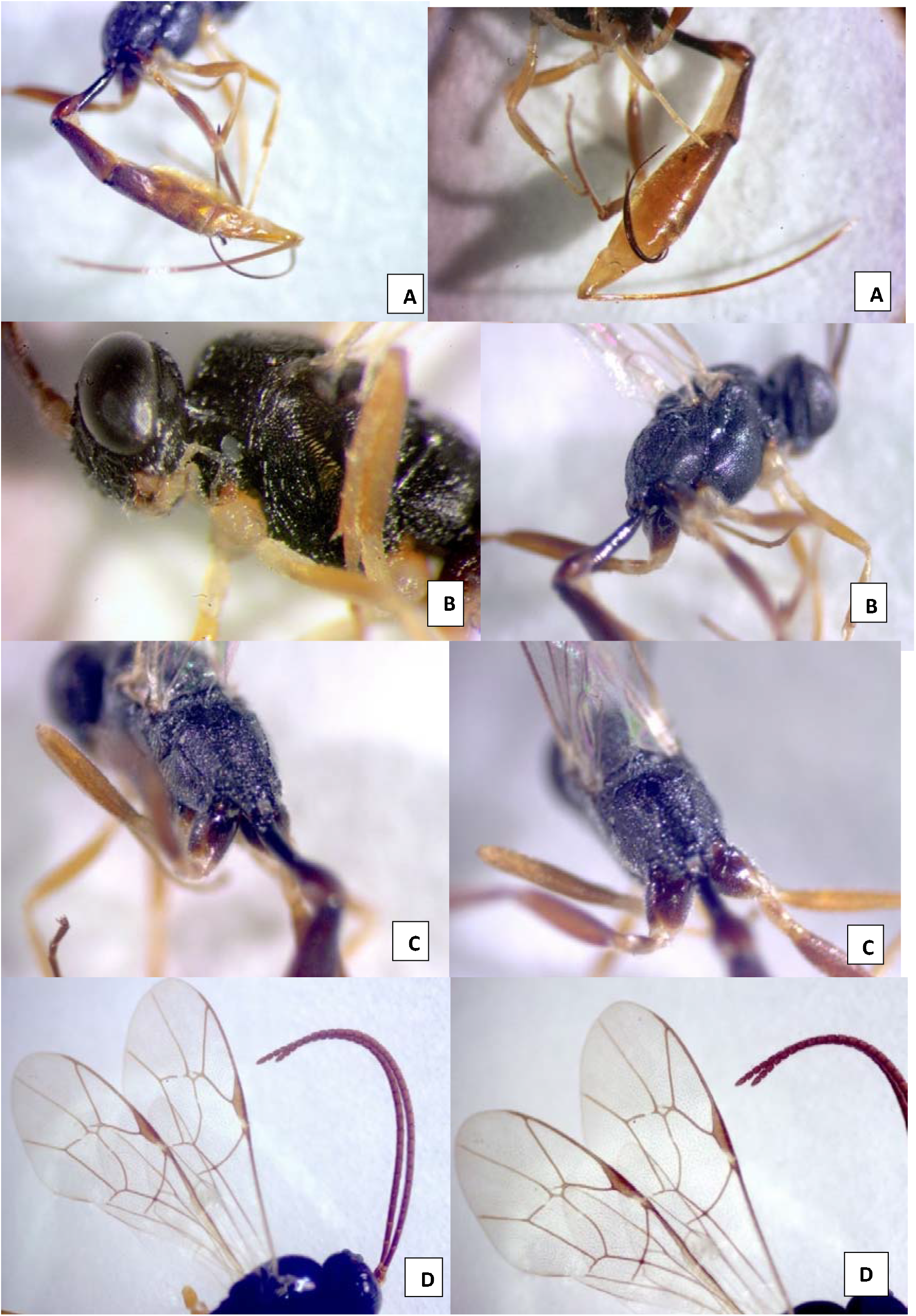
A. Metasoma reddish lateral view with reddish brown postpetiolus; B. Mesopleuron punctate; C. pentagonal elongated superomedial area; D. Antenna and Forewing, pterostigma brown, areolet not triangular.

## Discussion

*V. canescens*, accidently collected among parasitoids of the almond bark beetle *S. amygdali*, is a first record for Tunisia. The host is unknown, but the fruit crop where it was collected includes plum, apricot, peach and almond, which could host some harmful fruit lepidopterans.

*V. canescens* has been described under different names and assigned to different genera. The list of synonymies and generic combinations can be found in Yu et al. (2012).

*V. canescens* is a well-known parasitoid recorded on Pyralidae, Noctuidae, Tortricidae, Gelechiidae, Tineidae and Yponomeutidae for a total of Twenty-three host species (Van Noort 2022) (Fig. 5). It is often associated with pests of stored products with a worldwide distribution (Yu et al. 2012; Misfud et al. 2019; Vas 2019) (Fig. 6).

**Fig. 5.**
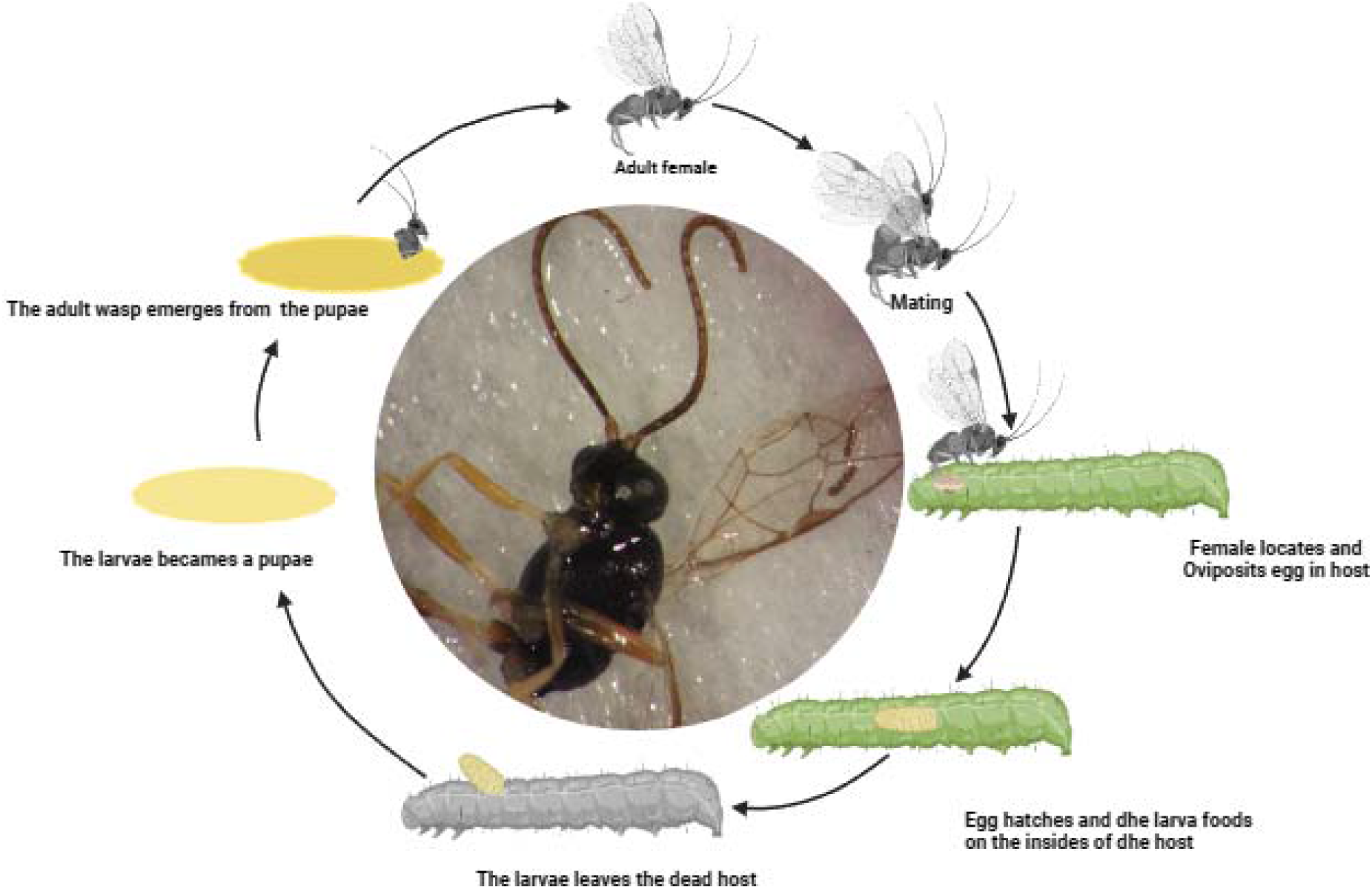
Life cycle of endoparasitoid wasps

**Fig. 6.**
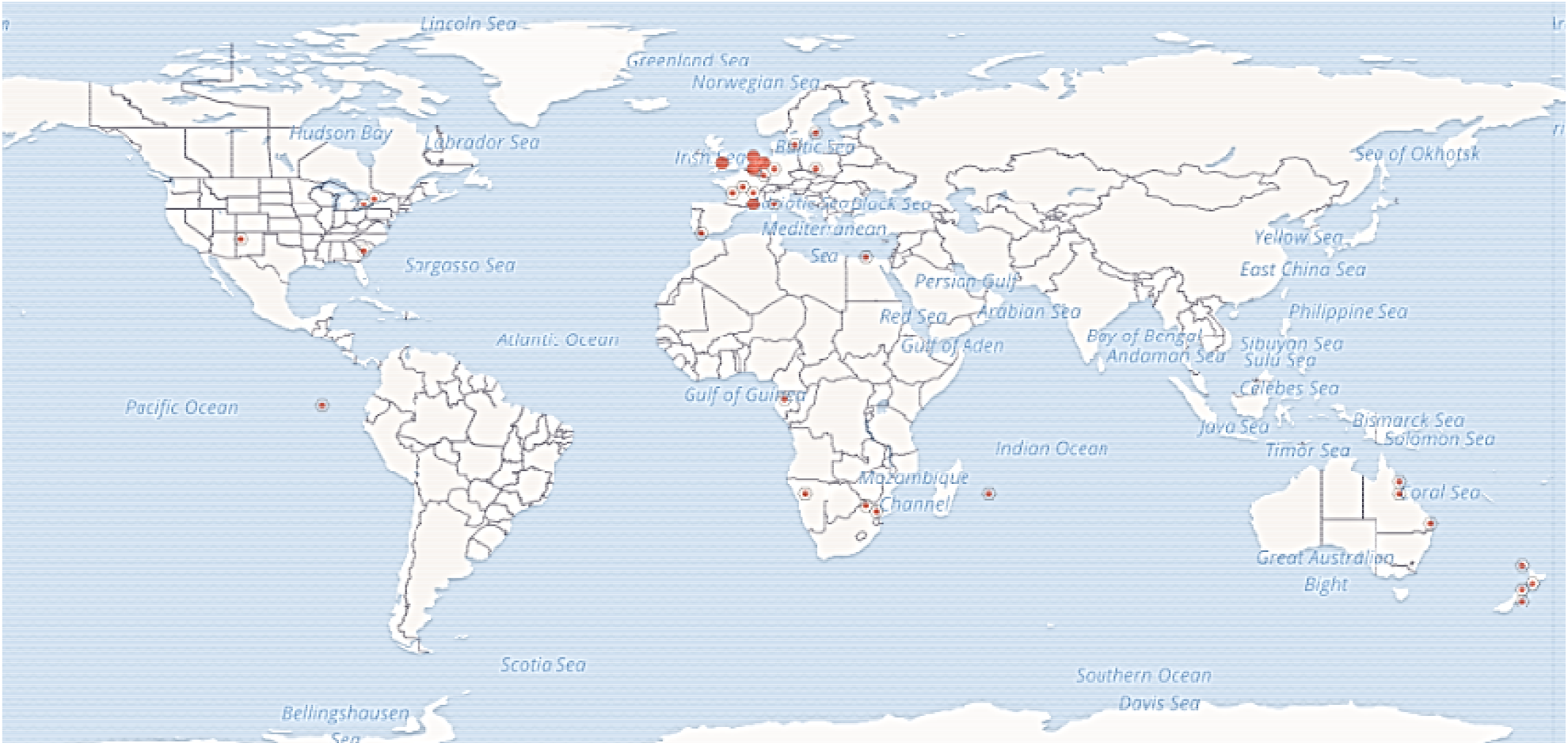
Distribution of *V. canescens* (Data from: *https://www.gbif.org/species/5028499; Venturia canescens (Gravenhorst, 1829) in GBIF Secretariat (2023). GBIF Backbone Taxonomy. Checklist dataset https://doi.org/10.15468/39omei accessed via GBIF.org on 2024-01-27*.)

*V. canescens* is a parasitoid of Twenty-three host species including Pyralidae, Noctuidae, Tortricidae, Gelechiidae, Tineidae and Yponomeutidae. In Tunisia, it was frequently associated with the carob moth *E*. ceratoniae, causing damages to dates (*P. dactylifera*), pomegranates (*P. granatum*), almonds (*P. dulcis*) and pistachio nut (*P. vera*) (Haouel et al. 2010; Braham 2015; Hached et al. 2020). The Professional Training Center of Agriculture in Tunisia where *V. canescens* was collected contains many fruit trees such as almond (*P. dulcis*), apricot (*Prunus armeniaca*), peach (*Prunus persica*), and plum (*Prunus domestica*) (Zeiri et al. 2015). It also contains pomegranates (*P. granatum*), pistachio nut (*Pistacia vera*), and citrus orchards. *V. Canescens* presence may be related to the presence of a mix of suitable hosts for *E. ceratoniae*. Superparasitism or laying eggs into parasitized hosts is a typical behaviour parasitoid such as *V. canescens* associated with man-made environments such as grain stores. The adult wasp uses its ovipositor to to deposit eggs in host larvae. *V. canescens* is a solitary koinobiont endoparasitoid, which means that a single egg is laid inside a host, from which will emerge, following a period of delayed development, a single adult. During this period the host continues to grow and feed (Jones et al. 2015).

## Conclusions

The identification of *V. canescens* was carried out using key for females of the genus *Venturia* to Western Palaearctic species (Vas 2020). and the specimen is reported as a first record in Tunisia. No male was collected, the host is unknown.

## List of Abbreviations

Not applicable

## Declarations

### Ethics approval and consent to participate

Not applicable.

### Consent for publication

Not applicable.

### Availability of data and materials

All data generated or analyzed during this study are included in the text.

### Competing interests

The author declare that they have no competing interests

## Acknowledgments

The Authors thank Dr. Mohammed Braham from “The Regional Center of Horticulture and Biological Agriculture” and Dr. Mohammed Braham from “The Olive Institute”, Sousse, Tunisia, for their help in the collection of data. I also thank Dr. Zoltán Vas, Hungarian Natural History Museum, Department of Zoology, Hymenoptera Collection for his help with the bibliography and the identification of the specimen. I am also thankful to Agrolab Italia and Agrolab Alimentalia for allowing me to use their stereoscopic microscope and for the help of the lab technician Vania Crestani with the photos. Special Thanks to Mr. Jeroen de Rond “Natural Media”, Mr. Senad Muric and Ms. Shahida Anusha Siddiqui for their valuable help and for English correction.

## Notes

### Competing Interest Statement

The authors have declared no competing interest.

